# Myeloarchitecture gradients in the human insula serve as blueprints for its diverse connectivity and function

**DOI:** 10.1101/839837

**Authors:** Jessica Royer, Casey Paquola, Sara Larivière, Reinder Vos de Wael, Shahin Tavakol, Alexander J. Lowe, Oualid Benkarim, Alan C. Evans, Danilo Bzdok, Jonathan Smallwood, Birgit Frauscher, Boris C. Bernhardt

## Abstract

Insular cortex is a core hub involved in multiple cognitive and socio-affective processes. Yet, the anatomical mechanisms that explain how it is involved in such a diverse array of functions remain incompletely understood. Here, we define a novel framework to test the hypothesis that changes in myeloarchitecture across the insular cortex explain how it can be involved in many different facets of cognitive function. Detailed intracortical profiling, performed across hundreds of insular locations on the basis of myelin-sensitive magnetic resonance imaging (MRI), was compressed into a lower-dimensional space uncovering principal axes of myeloarchitectonic variation. Leveraging two datasets with different high-resolution MRI contrasts, we obtained robust support for two principal dimensions of insular myeloarchitectonic differentiation *in vivo*, one running from ventral anterior to posterior banks and one radiating from dorsal anterior towards both ventral anterior and posterior subregions. Analyses of *post mortem* 3D histological data showed that the antero-posterior axis was mirrored in cytoarchitectural markers, even when controlling for anatomical landmarks and sulco-gyral folding. Resting-state functional connectomics in the same individuals and *ad hoc* meta-analyses showed that myelin gradients in the insula constrained affiliation to macroscale intrinsic functional systems, showing differential shifts in functional network embedding across each myelin-derived gradient. Collectively, our findings offer a novel approach to capture structure-function interactions of a key node of the limbic system, and suggest a multidimensional structural basis underlying the diverse functional roles of the insula.

## Introduction

Interfacing the frontal, temporal, and parietal lobes, the insular cortex is involved in a diverse set of functions ranging from lower-order sensorimotor processes to higher-order cognitive and socio-affective abilities (1, 2). Along with anterior cingulate and dorsolateral prefrontal cortices, the insula is a crucial hub of the ventral attention (or ‘salience’) network, and contributes to the prioritisation of sensory, visceral, autonomic, and attentional-executive processes guiding appropriate responses to relevant stimuli (3). Relatedly, many neuropsychiatric conditions involve perturbations of insular structure and function (4–6). This evidence establishes the insula as an important nexus that coordinates functions in a manner that is consistent with a core role across many cognitive domains (7). As such, through its contribution to multiple functional systems, the insula is a key structure to understand the principles shaping the coordination and competition of different networks in the human brain.

Current evidence suggests that the involvement of the insula in the diverse range of cognitive and affective features emerges from the complex connections it forms with other regions of the brain, and in particular through systematic changes between posterior and anterior regions (8, 9). Prior studies of subregional functional specialization in the insula have emphasized the presence of discrete functional clusters (10–12), as well as gradual changes in functional connectivity along its extent (13). Together, these studies highlight an architecture in which posterior regions of the insula are engaged primarily in sensory processing, while anterior regions are more connected to frontal areas involved in complex cognitive and affective states. Interestingly, this functional axis mirrors *post mortem* reports in animals and humans describing gradual cytoarchitectonic shifts from posterior to anterior banks of the insula (14–19). Although the changing microstructural context within its territory may indeed underlie the insula’s complex functional organization, the coupling between microstructural and functional architectures of this region remains to be examined in living humans.

Advances in high-field magnetic resonance imaging (MRI) allow classical neuroanatomical approaches to be extended to describe brain microstructure *in vivo*. In particular, contrasts sensitive to intracortical myelin content can quantify myeloarchitecture across the cortical mantle in single individuals and at the population level (20–23). With the development of multimodal co-registration techniques, microstructurally sensitive MRI data can be complemented with functional imaging methods, particularly resting-state functional MRI, to interrogate the interplay between structure and function in the living brain (24–26). Recent work has shown that maps of intracortical myeloarchitecture generated from myelin-sensitive contrasts appear to follow changes in task-based functional activation as well as intrinsic functional connectivity across the cortex (25, 27, 28). Quantitative intracortical *in vivo* profiling has also been leveraged to establish main axes of change in microstructural differentiation across the cortex in a data-driven manner, and has enriched our understanding of how microstructural composition may constrain the brain’s functional organization (24).

Our study interrogates how insula microstructure constrains its function by capitalizing on advanced analysis techniques tracking the correspondence between cortical microstructure and functional organization. Core to our approach was the application of a data-driven method revealing principal shifts in microstructural similarity, thus establishing a novel space representing differences in the similarity of intracortical myelin profiles. We explored how this new space mirrors shifts in cytoarchitecture, functional connectivity, and task-related activations across the insula, and assess the reproducibility of our findings across samples and with respect to distinct myelin-sensitive proxies. We identified two complementary dimensions of variation in insular myeloarchitecture, each with unique cellular properties and associated shifts in functional network embedding. Our findings offer an integrated empirical account of how the interplay between microstructure and function along multiple axes within the insular cortex gives rise to the unique role of this region in monitoring and modulating cortical information processing.

## Results

We studied 109 unrelated adults (62 women, 28.5±3.7 years) from the S900 release of the Human Connectome Project initiative (29, 30) that provides T1w/T2w images and cortical surface models. A digitized parcellation based on a classic cytoarchitectonic mapping of the human cerebral cortex developed by Von Economo and Koskinas was used to localize the insula, with boundaries slightly eroded to increase anatomical specificity (31). An equivolumetric transformation (32) generated a series of intracortical surfaces used to sample T1w/T2w intensities across cortical depths, yielding rich microstructural profiles at each insular vertex (**Figure 1A**). Following a recent approach (24), we performed pairwise correlations of the microstructural profiles across vertices, covaried for average intensity profile, to generate a matrix for each participant that captured microstructural profile similarities. Matrices were averaged across participants, and the result was transformed into an affinity matrix (**Figure 1B**). This affinity matrix was mapped onto a lower dimensional manifold using openly available non-linear techniques aimed at finding hidden patterns of dominant variation (33), similar to recent work that identified functional connectivity gradients based on resting-state fMRI (34, 35).

**Figure 1.**
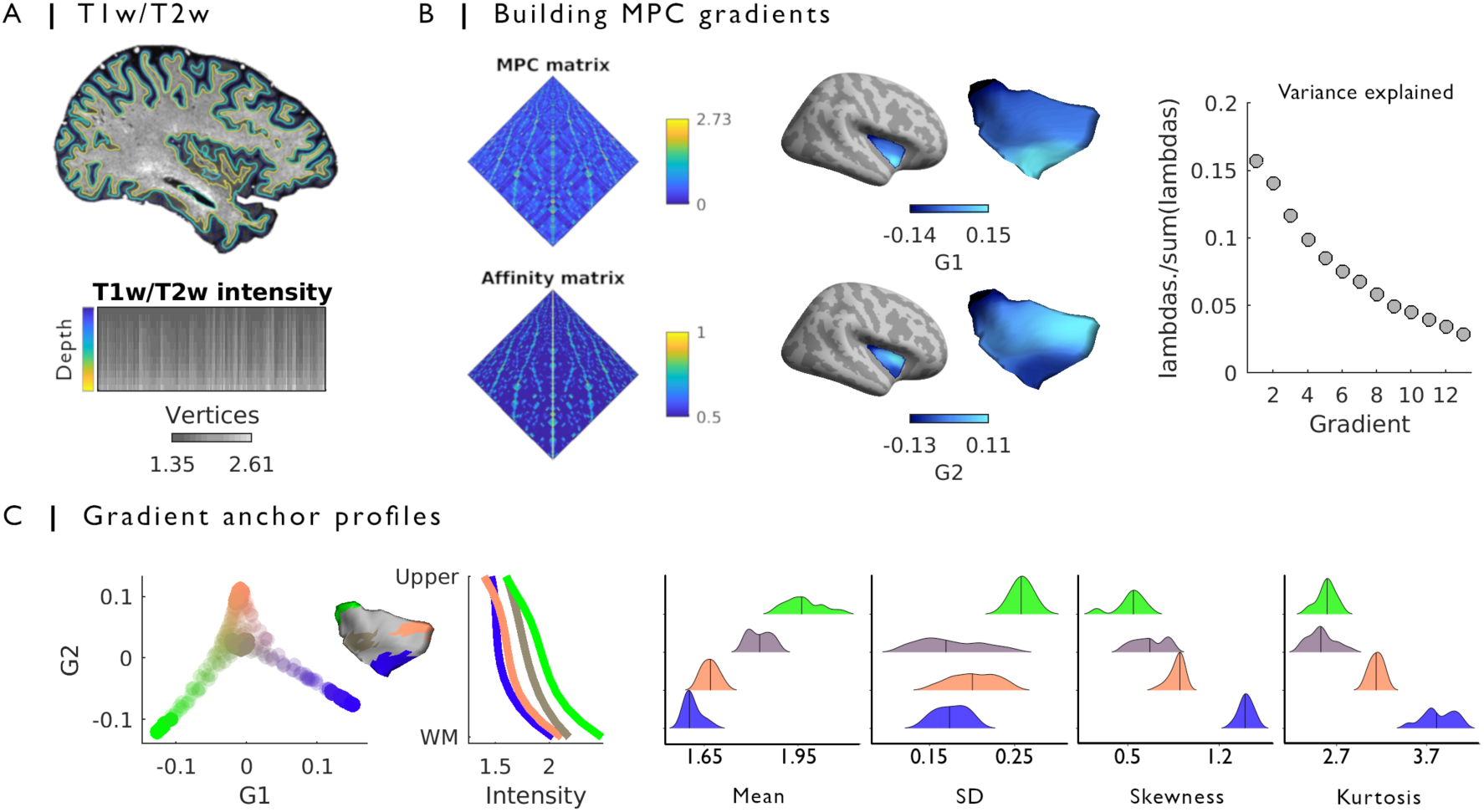
In vivo microstructural gradients of the insular cortex. **(A)** T1w/T2w intensities were sampled from the pial to the white matter boundary in each participant, yielding vertex-wise *in vivo* microstructural profiles. **(B)** We applied microstructural profile covariance (MPC) to the intensity profiles of vertices within the insula of each participant. The group-averaged MPC matrix was transformed into an affinity matrix, reflecting similarity in microstructure. This matrix was subjected to nonlinear dimensionality reduction to identify principal axes of microstructural differentiation. The principal gradient (G1) depicted a transition from posterior to ventral anterior regions, while the second gradient (G2) differentiated dorsal anterior and both posterior and ventral anterior areas. **(C)** Four highly differentiated compartments emerged from this 2D manifold (coloured dots and areas projected onto the insula surface). Line plot shows average T1w/T2w intensity profiles in selected compartments. Ridge plots show statistical moments of compartment intensity profiles, confirming gradual changes in depth-specific T1w/T2w profiles.

### Principal gradients in microstructural similarity within the insular cortex

The principal gradient (G1) of insular microstructure profile similarity depicted a transition from posterior to ventral anterior regions, while the second gradient (G2) radiated away from posterior and ventral anterior areas towards dorsal anterior regions (**Figure 1B**). Both gradients explained 29.9% of variance in T1w/T2w profile similarity across the right insula (see **Figure S1** for equivalent results in the left hemisphere). To illustrate intracortical mechanisms driving G1 and G2, we selected four distinct compartments in the manifold spanned by both gradients. Compartments were highly differentiated with respect to their overall profiles, particularly in mid- and deeper cortical layers (**Figure 1C**). Differences in microstructure were examined using statistical moment theoretical parameterization (*i.e.*, by calculating mean, standard deviation, skewness, and kurtosis of all intracortical T1w/T2w profiles), an approach previously used to guide boundary definition in cytoarchitectonic studies of *post mortem* data (36, 37). This highlighted large differences between ventral anterior and posterior insular gradient anchors. Interestingly, dorsal anterior and medial compartments were located between more agranular ventral anterior and granular posterior subregions, mirroring findings of animal cytoarchitectonic studies (14–16) showing the predominance of dysgranular cortex in this region.

### Cytoarchitectonic underpinnings of microstructural gradients

We next examined the relationship of these gradients to gold standard measures of microstructure, by assessing the cytoarchitectonic underpinnings of G1 and G2 with BigBrain (**Figure 2A**), a 3D histological reconstruction of a Merker-stained *post mortem* human brain (38). BigBrain intensity profiles, reflecting soma size and cellular density, were explored within the four compartments defined previously. While ventral anterior and medial regions presented with higher intensity in deeper cortical layers, dorsal anterior and posterior areas showed an opposite pattern with progressive intensity decreases with cortical depth. We next explored whether the gradients were correlated with depth-wise intensities. Correlations between histological intensity and G1 (mean Spearman r=0.21, SD=0.20) were stronger in deeper cortical layers, while correlations with G2 (mean Spearman r=-0.05, SD=0.06) remained weak at all cortical depths. Notably, strongest histological correlations with G1 were obtained at 87.5% depth (Spearman r=0.51; **Figure 2B**). Higher intensity values were seen in the anterior insula, particularly the ventral anterior region, possibly reflecting the presence of large pyramidal neurons located largely in deeper cortical layers of this area (16, 39, 40). These associations were robust to variations in cortical morphology, specifically cortical thickness and curvature, highlighting the microstructural specificity of the association between deep layer histological intensity and the principal myeloarchitectonic gradient (Spearman r=0.44; **Figure 2C**). To determine the significance of the above correlations, we carried out non-parametric tests that account for shared spatial auto-correlation in gradient and histological data (41). This approach showed that correlations between G1 and deep layer histological intensity remained consistent (controlled for thickness and curvature: *p*_spatial_=0.05; raw: *p*_spatial_=0.01). Thus, our findings indicate that cytoarchitecture, and in particular cortical depth-specific changes in cell density and size, underpin the principal microstructural gradient differentiating ventral anterior and posterior insular subregions.

**Figure 2.**
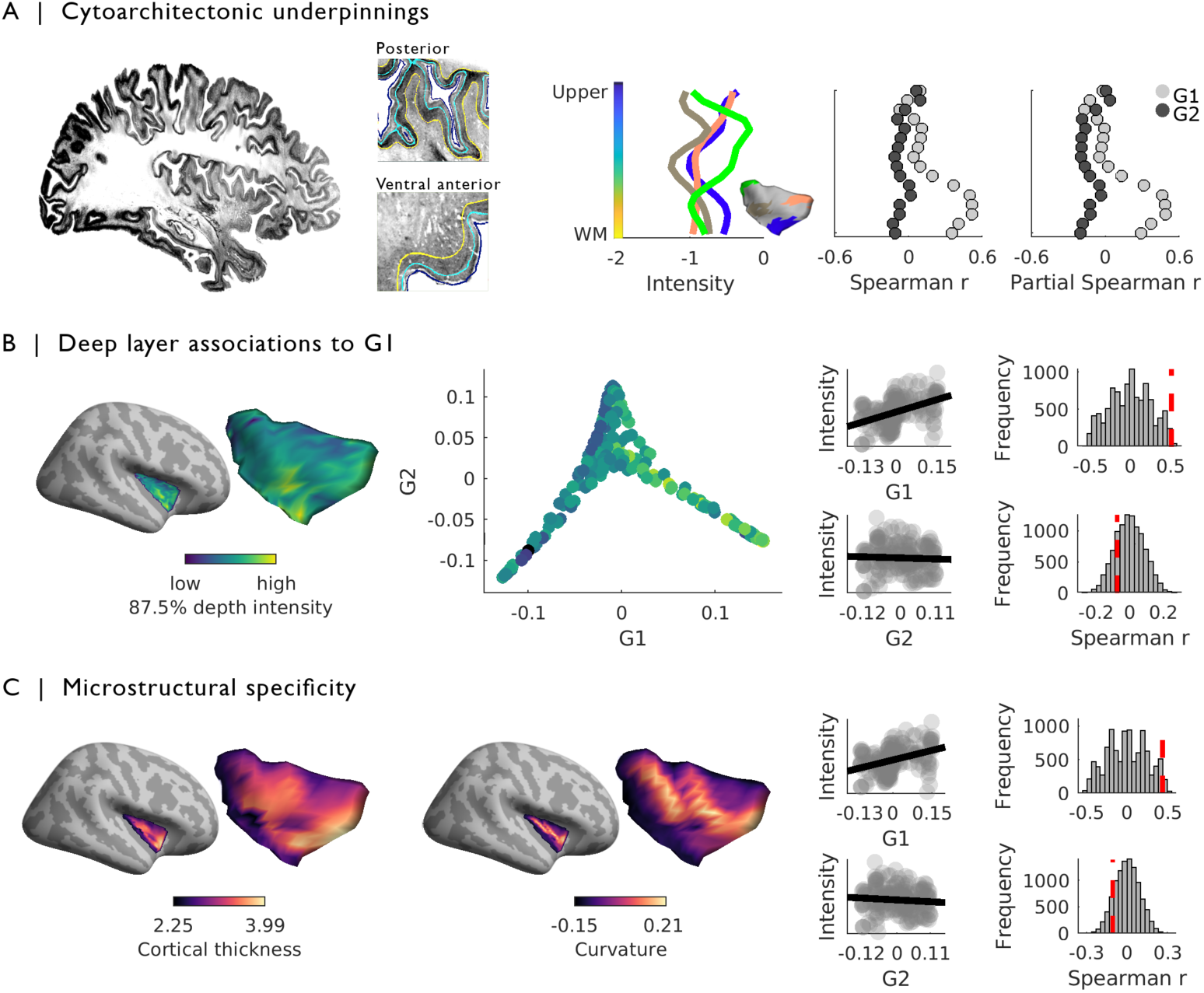
Cytoarchitectonic underpinnings of in vivo microstructural gradients in the insula. **(A)** Based on BigBrain, intracortical histological intensity was sampled along 16 equivolumetric surfaces, yielding distinct intensity profiles at each insular vertex. Comparing average intensity profiles across previously described gradient compartments showed prominent differences in histological intensity between ventral anterior and posterior insular banks in deeper cortical layers. Depth-specific rank correlations between histological intensities and each of the MPC gradients showed that stronger correlations were restricted to deeper cortical layers along G1, both when using raw intensity values and when controlling for cortical thickness and curvature. **(B)** Histological intensity at 87.5% depth was significantly correlated with G1, but not G2, **(C)** and findings at this depth were robust to variations in cortical thickness and curvature. Correlations with raw and residual deep-layer intensity (controlling for cortical morphology) were robust to null model testing accounting for shared spatial auto-correlation in gradient and histological data.

### Associations to macroscale functional connectivity

Having documented distinct gradients within insular microstructure, we next examined whether they have any associations with observed neural function as described by functional connectivity at rest. Changes in whole-cortex functional connectivity profiles along both microstructural gradients were examined in relation to established functional communities (**Figure 3A**) (42). Moving from posterior to ventral anterior insular regions along G1, connectivity fingerprints shifted from a preferential affiliation to somatomotor and visual networks, towards a pattern containing both unimodal and ventral attention (salience) networks. In contrast, moving along G2 from ventral anterior and posterior regions towards dorsal anterior insula, we observed a shift from a dominant unimodal network fingerprint towards increasing participation in dorsal and ventral attention networks, culminating in the transmodal frontoparietal network at the most dorsal extent of this trajectory. Our results demonstrate diverging transitions in functional connectome embedding along the changing microstructural landscape of the insula, with changing affiliation from unimodal to attention and transmodal regions following the main axes of myeloarchitectonic differentiation along this structure.

**Figure 3.**
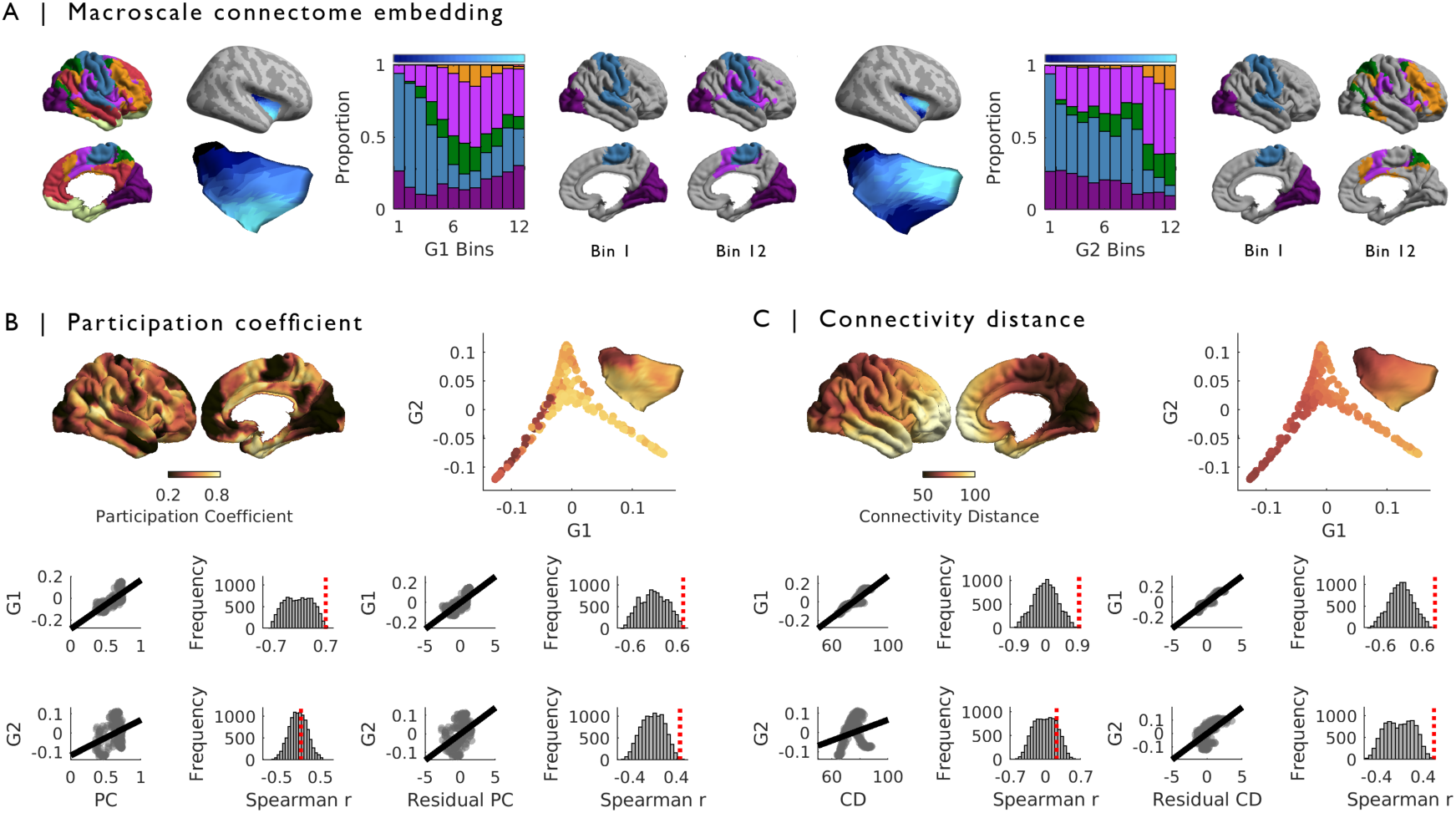
Microstructural gradients track functional networks architectures. **(A)** Affiliation to established functional communities (42) was assessed by calculating the proportion strongly connected nodes (>10%) falling within the boundaries of each community. Functional connectivity patterns of G1 (left) and G2 (right) bins were computed separately. Functional communities with proportions above chance level (1/7 networks = 0.14) for anchor bins are displayed on the cortical surface. **(B)** Vertex-wise participation coefficients (PC) were highly correlated with G1 (top row). G2 was not significantly correlated with raw PC values, but a significant correlation was obtained with standardized residual PC values obtained from a simple linear model of PC predicted by G1 (bottom row). **(C)** Vertex-wise connectivity distance (CD) was highly correlated with G1 (top row). G2 was not significantly correlated with raw CD values, but we observed a significant correlation between G2 and standardized residual CD values obtained from a simple linear model of CD predicted by G1 (bottom row).

Next, graph theoretical analyses were applied to quantify the manner through which graded changes in insular microstructure vary with connectome wide organization of neural function. First, we identified that the participation coefficient (PC; **Figure 3B**), a measure capturing the heterogeneity of rs-fMRI connectivity (see *Materials and Methods*), increased strongly along G1 (Spearman r=0.69; *p*_*spatial*_<0.01) but not along G2 (Spearman r=0.06; *p*_*spatial*_=0.8). However, the correlation between PC and G2 increased when controlling PC for G1 (Spearman r=0.46; *p*_*spatial*_<0.01), suggesting that the dorsal-ventral axis of microstructural similarity described by G2 also partly supports functional diversity in the insular cortex. The association between G1 and PC was robust to changes in G2 values (Spearman r=0.75; *p*_*spatial*_<0.01). Moreover, we observed corresponding changes in functional connectivity distances (CD; **Figure 3C**) using a recently developed metric that captures the average geodesic distance of a node’s connectivity profile (35, 43). Indeed, connectivity distance was predicted by increases in G1 (Spearman r=0.95, *p*_*spatial*_<0.01), but much less so by G2 (Spearman r=0.22; *p*_*spatial*_=0.37). Similar to what was observed with PC, controlling CD for changes in G1 revealed a stronger association between CD and G2 (Spearman r=0.58; *p*_*spatial*_<0.01), while controlling for G2 in the association between G1 and CD had little effect (Spearman r=0.97, *p*_*spatial*_<0.01). These results suggest a more local influence on neural function in the posterior insular, while ventral and dorsal anterior regions affiliated with attentional-executive networks were associated with organisation of cortical function at a broader scale. Together, analysis of the relationship with functional connectivity establish that insula gradients show unique associations both in terms of local and long-range patterns of connectivity, and their associations with neural systems that serve distinct functions.

### Differential associations to cognitive domains

Our study has established that overlapping patterns of myeloarchitetonic transitions within the human insula interact with other regions in a distributed and unique manner. Our final analysis examines whether these unique patterns can account for the range of functions that the insula has been implicated with. To establish correspondence to cognitive taxonomies, we computed spatial associations between the microstructural gradients and local insular activations derived from meta-analytic term maps (24, 34, 44) (**Figure 4A**). Changes in term associations along G1 reflected a transition from sensory mechanisms in the posterior insular anchor, and culminated in cognition and emotion-related terms in anterior regions. A similar approach was carried for G2, which discriminates between ventral and dorsal anterior segments (**Figure 4B**). This highlighted a shift from emotion-related terms mapping onto ventral anterior regions, towards primarily attentional-executive affiliations in dorsal anterior insular banks (10, 11, 45). These findings emphasize the contributions of each microstructural gradient to the existence of local functional hierarchies within the insular cortex. Indeed, the principal microstructural gradient, which we found to be grounded in insular cytoarchitecture, related to functional shifts evolving from sensory to affective processes. Conversely, the dorsal-ventral axis of microstructural differentiation reflected by G2 dissociated distinct aspects of higher-order processing in which the insula is implicated, specifically affective and attentional-executive mechanisms.

**Figure 4.**
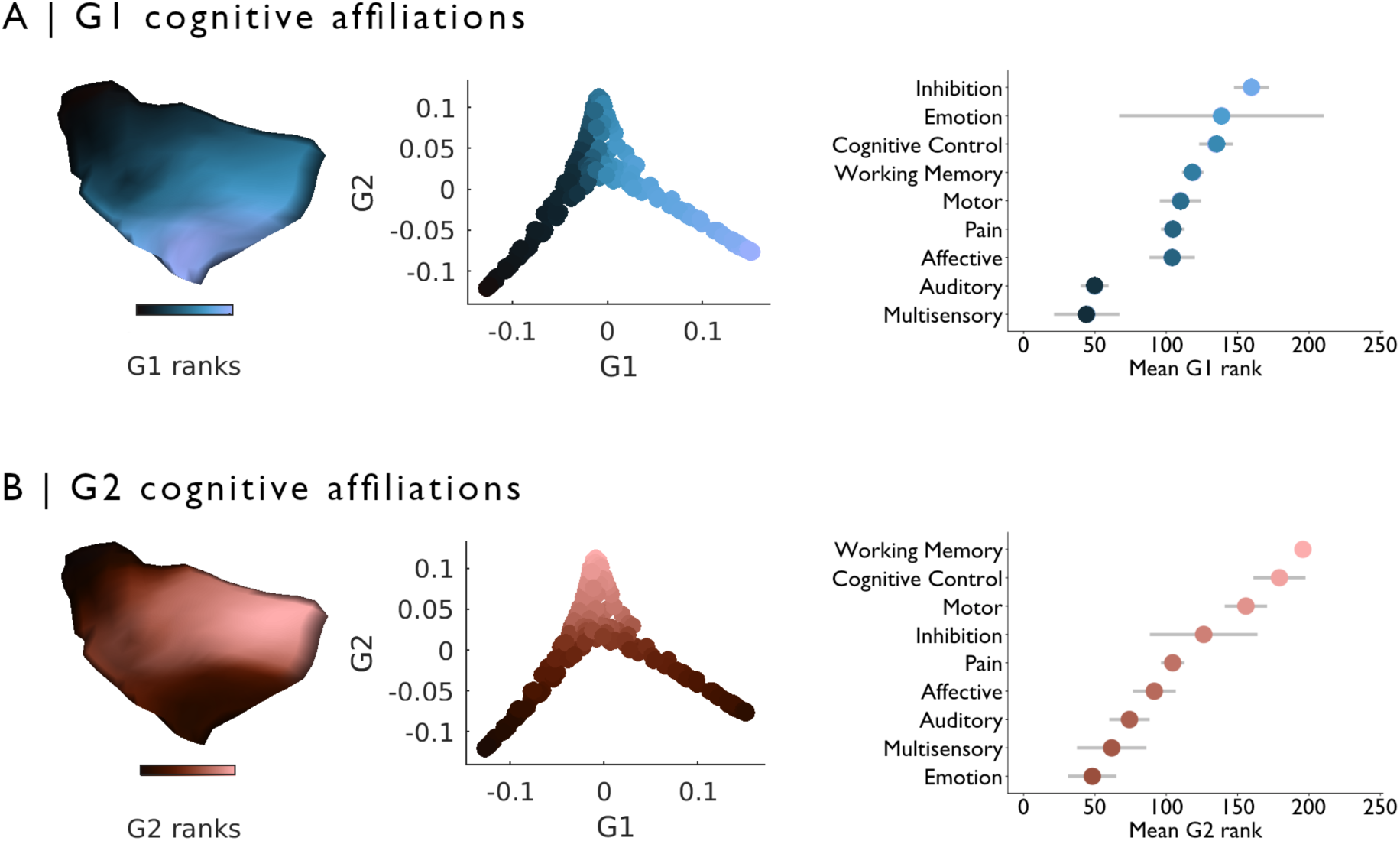
Insular cortex microstructural gradients are linked to distinct cognitive taxonomies. **(A)** Meta-analytic term maps derived from Neurosynth revealed shifts in local insular affiliations along G1. Mean G1 rank within each term activation map showed a transition from unimodal sensory processes in the posterior anchor, towards cognitive and affective mechanism in the anterior anchor. **(B)** Shifts along G2 highlighted a divergence between dorsal anterior and ventral anterior anchors, and corresponding term activation maps shifted from primarily affective affiliations in ventral anterior regions, towards higher-order attentional-executive terms preferentially mapping to dorsal anterior areas. Error bars denote 95% CIs.

### Sensitivity analyses

Several analyses assessed robustness of the discovered microstructural axes of differentiation within the insula. First, analysing the left insula revealed virtually identical microstructural gradients to those of the right hemisphere, indicating inter-hemispheric consistency (**Figure S1**). Gradients generated in each hemisphere furthermore accounted for a comparable microstructural variance (left: 30.2%, right: 30.4%). Findings were also consistent with respect to different matrix thresholding procedures (**Figure S2**), and robust when alternative dimensionality reduction techniques were used (**Figure S3**). Results furthermore generalized to an independent cohort of 34 healthy adults (14 women; 29.38±6.04 years) scanned at our site using a different myelin sensitive contrast, namely quantitative T1 relaxometry (MICA-MTL cohort; see *Materials and Methods*). Gradients in this replication cohort were indeed highly similar to the original findings (Spearman correlations in right/left: G1: r=0.98/0.99; G2: r=0.97/0.99; *p*<0.001; **Figure S4**).

## Discussion

Using novel imaging methods and advanced analysis techniques, our work highlighted two complementary dimensions of microstructural differentiation in the human insula, each with distinct cellular underpinnings and functional affiliations. Through a systematic profiling of intracortical T1w/T2w image intensities, a proxy of cortical myelination (20), our findings provided the first translation of seminal histological observations to myeloarchitectonic investigations of the insula in living humans. We demonstrated that shifts in insular myeloarchitecture follow two major gradients, one evolving in anterior-to-posterior (G1) and one in dorsal-to-ventral direction (G2). Leveraging *post mortem* 3D histological reconstructions, we found that G1 was linked to regional cytoarchitecture, specifically soma size and density in deeper cortical layers, which could not be explained by changes in mesoscopic cortical morphology and sulco-gyral landmarks. These gradients were also differentiable based on their associations with connectome-wide patterns of neural organisation. While G1 anchors were both affiliated to unimodal cortices with an additional contribution of the salience network in the ventral anterior anchor, connectivity changes along G2 reflected a transition from unimodal and salience networks to attention and transmodal systems. Moreover, the microstructural gradients mapped local functional hierarchies within the insular cortex. The cytoarchitectonically grounded G1 supported functional shifts from sensory to affective processes, while the dorsal-ventral axis of microstructural differentiation reflected by G2 dissociated distinct aspects of higher-order processing, specifically affective and attentional-executive mechanisms. Importantly, our microstructural gradient findings were consistent in both hemispheres, robust against algorithm variations, and replicable across two independent samples with different myelin MRI contrasts. Collectively, our findings provide robust *in vivo* evidence for multiple graded changes in insula myeloarchitecture that ultimately contribute to its heterogeneous neural functional profiles in humans.

Our work uncovered a close correspondence between the principal microstructural gradient in the human insula derived from *in vivo* MRI and descriptions of its microstructural heterogeneity from prior histological work in cadaver brains. These classical neuroanatomical investigations performed in the first half of the 20^th^ century subdivided the insula into distinct cytoarchitectonic zones, ranging from only 2 clusters (18, 46) up to approximately 30 subareas (47, 48), increasingly emphasizing graded shifts in laminar differentiation forming concentric rings along the ventral anterior to posterior insular bank (14–16). In line with these observations, we found largest myeloarchitectonic differences between ventral anterior and posterior insular regions. These areas served as the anchors of our principal microstructure gradient, while the dorsal anterior area was positioned as a transition zone in this axis. Harnessing a 3D histological reconstruction of a *post mortem* human brain, we could establish specificity of our findings to intracortical microstructure, by demonstrating correspondence between the main axis of myeloarchitectonic differentiation and the size and density of cell bodies in deeper cortical layers. Interestingly, this finding was mainly driven by higher histological intensities in deeper cortical layers of anterior insular banks, and particularly the ventral anterior insula. This finding may reflect the presence of a previously observed subtype of pyramidal neurons supporting long-range communication to other areas within the anterior insula (39, 40, 49). Crucially, the cytoarchitectonic underpinnings of the principal myeloarchitectonic gradient were robust to variations in cortical thickness and curvature. This result is compatible with accounts suggesting little correspondence between microstructure and morphological properties, such as gyral boundaries and sulcation within the insula (15, 16, 50), and suggests specificity of our findings for the intracortical microstructural context.

Previous work in the macaque has shown that the balance of infragranular to supragranular projections within a given cortical area offers a possible microcircuit basis of its functional role. Indeed, the laminar origin of cortical projections are thought to relay information transit along the cortical functional hierarchy, such that sensory areas conveying feedforward information to higher-order cortical structures project mostly from supragranular layers, while paralimbic regions involved in feedback processing predominantly project from infragranular layers (51–53). Our systematic analysis of local T1w/T2w image intensity profiles via statistical moment parameterization, an approach applied in several histological studies (36, 37), indicated gradual changes in the shape of intracortical myelin profiles from anterior to posterior insular regions. These findings may thus reflect transitions in the laminar origin of projections seen along G1, a finding also in line with the BigBrain results showing higher clustering of cell bodies in deeper cortical layers in the agranular ventral anterior insular area compared to the posterior anchor. In the future it may be possible to fully understand the relationship between neural activity in the insula and the underlying microcircuitry using computational models as well as neural recording and stimulation paradigms (54).

More generally, our study suggests that overlapping myeloarchitectonic gradients within the insula offer an explanation for the observation that neural signals within this regions are associated with a wide range of cognitive and affective processes. Previous work has highlighted functional connectivity gradients (13) along an anterior-posterior axis of the insula, mirroring the principal microstructural gradient identified here. Importantly, our study builds on this finding by demonstrating that there are multiple graded patterns within insular cortex, each with unique neural features and that have significant associations with observed patterns of functional connectivity. We found that the principal gradient G1 was focused on local cortical features, linked to cell density and dissociated unimodal networks in posterior insula from attentional systems in anterior regions. In contrast, functional connectivity shifts along G2 were linked to longer distance connectivity patterns, and in particular highlighted a distinction between unimodal and transmodal systems seen in a prior study (Margulies et al., 2016). Our findings, therefore, emphasize that the insula is organized along multiple axes of microstructural differentiation that each reflect a different balance of local and distal communication, and are associated with neural systems that support qualitatively different neural functions. This microstructural organisation of the insula allows patterns in this region to reflect multiple features of the organisation of the cortex as a whole (and vice versa), which may explain how neural signals in the insula can convey information regarding the wide range of functional states. In this way our study establishes that multiple overlapping graded patterns within the microstructural architecture within the insula as an important candidate mechanism for why this region plays a pivotal role in such a wide range of cognitive and effective states.

We close by highlighting the importance of open science initiatives, such as the Human Connectome Project, to provide targeted studies on the coupling of healthy brain structure and function. The increasing importance allocated to reproducibility and generalizability of findings in the neuroimaging literature promises to lead to a more robust understanding of brain organization. In the present work, results were found to be consistent across different myelin-sensitive MRI sequences and scanning sites, and all insular gradient data and code has been made openly available for independent verification of our conclusions. The robustness of our results across healthy cohorts is encouraging for future studies aiming to clarify the role on the insula in disease. Indeed, in light of the high susceptibility of insular microstructure and function to a range of neurodevelopmental and neurodegenerative conditions (55, 56), we hope that the microstructurally-governed microcircuit layout of the insula identified in this work may guide future structure-function studies on this intriguing structure in diseased populations. Our findings provide a framework to address coupled perturbations in insula microcircuity, macroscale function, and clinical phenomenology, and promise to refine our understanding of the many brain disorders that present with considerable structural and functional anomalies of the insular cortex.

## Materials & Methods

### Sample characteristics

We studied 109 unrelated adults (62 women, 28.5±3.7 years) from the S900 release of the Human Connectome Project initiative, HCP (29, 30), in whom T1w/T2w images, cortical surface models, and four resting-state functional scans were available. Participants from this initiative, which served as our Discovery sample, provided informed consent for open sharing of their deidentified data, approved by the Washington University Institutional Review Board as part of the HCP.

### MRI acquisition

Structural and functional scans were acquired on the HCP’s custom 3T Siemens Skyra with a 32-channel head coil. Two T1w scans with identical parameters were acquired with a 3D-MPRAGE sequence (0.7mm isotropic voxels, matrix=320×320, 256 sagittal slices, TR=2400ms, TE=2.14ms, TI=1000ms, flip angle=8°; iPAT=2). In addition, two T2w scans were acquired using a 3D T2-SPACE sequence and with identical geometry to the T1w images (TR=3200ms, TE=565ms, variable flip angle; iPAT=2). Both T1w and T2w scans were acquired on the same day. Four resting-state functional scans were acquired with multi-band accelerated 2D-BOLD echo-planar imaging (2mm isotropic voxels, matrix=104×90, 72 sagittal slices, TR=720ms, TE=33ms, flip angle=52°; mb factor=8; 1200 volumes/scan). A fixation cross was presented on screen, and participants were asked to fixate the cross, and not to fall asleep. The four functional scans were split over 2 days, with 2 scans acquired per day.

### Image preprocessing

Structural and functional MRI data underwent HCP’s minimal preprocessing (30, 57, 58), with minor adjustments to incorporate both T1w and T2w (20). Following intensity nonuniformity correction, T1w images were divided by aligned T2w images to produce a single volumetric T1w/T2w image per subject (20). As for resting-state fMRI, timeseries were first corrected for gradient nonlinearities and head motion. The R-L/L-R blipped scan pairs were used to correct for geometric distortions. Distortion-corrected images were warped to the same space as the T1w images using rigid body and boundary-based linear registrations (59). These transformations were concatenated with the transformation from native T1w to MNI152 to warp functional images to MNI152. Further processing involved removal of the bias field (as calculated for the structural image), brain extraction, and whole-brain intensity normalisation. A high-pass filter corrected timeseries for drifts. Additional noise was removed using ICA-FIX (58). Tissue-specific signal regression was not performed (60, 61).

### Localizing the insula

Conte69 templates were downsampled to the fsaverage5 template by retaining data from the nearest neighbouring vertex as defined by their Euclidean distance. The insula was localized within this space by applying a slightly eroded mask (Gaussian smooth; FWHM=4) derived from a digitized parcellation of Von Economo and Koskinas’ cytoarchitectonic mapping of the human cerebral cortex (31).

### Microstructural profile covariance (MPC) and gradient mapping

#### Surface construction and sampling

We generated 14 equivolumetric intracortical surfaces (32) that sampled T1w/T2w intensities across cortical depths, yielding distinct intensity profiles reflecting microstructural composition of each vertex contained within the insular boundaries. This number was selected based on the results of stability analyses of the MPC matrix (see *below*) performed in previous work (24). Vertex-wise microstructural profiles covering the pial to white matter boundary were estimated across these surfaces for both the left and right insula (**Figure 1A, S1**). Data sampled from surfaces closest to the pial and white matter boundaries were discarded to reduce partial volume effects. Vertex-wise intensity profiles were smoothed with a Gaussian kernel (FWHM=3) prior to building the MPC matrix using tools from SurfStat for Matlab (http://mica-mni.github.io/surfstat) (62).

#### MPC construction

We cross-correlated microstructural profiles across all insular vertices to generate a matrix for each participant that captured microstructural profile correlations. The MPC approach yielded similarity matrices in myelin proxies across the insula, while controlling for average intensity. Only positive correlations were retained, and values were log-transformed (**Figure 1B**). This procedure was performed separately for the left and right insula. Insula-wide MPC matrices for each hemisphere were averaged across participants for further analyses.

#### Gradient mapping

Microstructural gradient in the insula were identified with the BrainSpace toolbox for Matlab (https://github.com/MICA-MNI/BrainSpace) (63). Group-averaged MPC matrices were thresholded row-wise to retain only the top 10% of values as in prior work (24, 25, 34, 35) and subsequently transformed into normalized angle affinities (**Figure 1B**). Affinity matrices were projected into a lower dimensional manifold using diffusion map embedding (33). This nonlinear technique identified principal gradient components explaining Eigenvalues in the MPC matrices in descending order. As in previous studies (24, 25, 34, 35), we set the α parameter of this algorithm to 0.5, a decision that has been suggested to retain the global relations between data points in the embedded space and seems robust to noise in the covariance matrix. Gradients were displayed in the embedding space, or mapped to the pial surface of the fsaverage5 template and visualized via SurfStat (62).

#### MPC gradient characterization

Variations in T1w/T2w intensity profiles along the space composed of the first two microstructural gradients were characterized using statistical moment parameterization (36, 37, 64). We calculated the mean, standard deviation, skewness, and kurtosis of intracortical T1w/T2w profiles of each insular vertex. Selected vertices composing G1 and G2 anchors and an intermediate transition zone were grouped into four distinct compartments. Inclusion of vertices in each compartment was defined using histogram bin edges of G1 and G2 obtained from 12 discrete bins containing an equal number of vertices, and the distribution of values of all vertices in each compartment was visualized using ridge plots (**Figure 1C**).

### Cytoarchitectonic markers

#### Histological data acquisition and preprocessing

A high-resolution Merker-stained 3D volumetric histological reconstruction of a *post mortem* human brain from a 65-year-old male was obtained from the open-access BigBrain repository (https://bigbrain.loris.ca/main.php) (38). The brain was embedded in paraffin, sliced in 7,400 20µm coronal sections, silver-stained for cell bodies (65), and digitized. Resulting images were manually inspected for artefacts, and automatically repaired by aligning the images to *post mortem* MRI, applying intensity normalization, and block averaging (66). Geometric meshes approximating the outer and inner cortical interface (*i.e.*, the GM/CSF boundary and the GM/WM boundary) with 163,842 matched vertices per hemisphere were also available (67). Analyses were performed on inverted images, on which higher staining intensity reflects higher cellular density and soma size, and data were downsampled to 100 µm for computational efficiency.

#### Surface construction and sampling

We generated 16 intracortical surface-based intensity profiles from the histological data using a similar procedure to the previously described T1w/T2w intensity profiling. (**Figure 2A**). Staining intensity values, reflecting soma size and cell density, were systematically sampled along these surfaces, yielding distinct intensity profiles at every cortical vertex. Vertex-wise histological intensity profiles were downsampled to fsaverage5, retaining data from the nearest neighbouring vertex. Standardized residuals from a simple linear model of surface-wide intensity values predicted by the pial *y* coordinate were used in further analyses to account for an anterior–posterior increase in intensity values across the BigBrain due to coronal slicing and reconstruction (38).

#### Histological infragranular intensity

Intensity values of vertices within the previously defined insula mask were correlated with G1 and G2 across all cortical depths (**Figure 2B**), showing stronger rank correlations with the principal gradient in deeper cortical surfaces. Strongest correlations with G1 were seen at 87.5% depth, and values extracted from this surface were retained for further analyses (**Figure 2C**). The association between histological intensity at this depth and both gradients was also assessed when controlling for cortical morphological features. Individual estimates of cortical thickness and mean curvature were provided by HCP as part of their minimal processing pipeline (30), obtained using FreeSurfer 5.3.0-HCP (68–70). Standardized residuals from a simple linear model of histological measures predicted by cortical morphological features were related to G1 using Spearman rank correlations. Statistical significance of reported correlations was assessed with Moran spectral randomization (41) implemented in the BrainSpace toolbox (63). This method computes a metric for spatial auto-correlation, Moran’s *I*, and generates normally distributed data with similar auto-correlation. Moran eigenvector maps were computed based on each insula vertex’s direct neighbouring nodes on the fsaverage5 pial surface. Spectral randomization implemented a singleton procedure (10,000 permutations).

### Functional affiliation of microstructural gradients

#### Intrinsic functional connectomics

Surface-mapped resting-state functional MRI timeseries were downsampled to the fsaverage5 template, and timeseries averaged within the previously defined G1 and G2 histogram bins. We computed the correlation between the average time series of each microstructural gradient bin and the time series of all other cortical vertices for each participant. Connectivity matrices underwent Fisher R-to-Z transformations, and were averaged across participants. Matrices were thresholded to retain the top 10% of connections of each gradient bin, as in prior work (34). A community parcellation of the cortex (42), provided on fsaverage5 (https://github.com/ThomasYeoLab/), was used to quantify the affiliation of each gradient bin to macroscale functional communities. Changes in the proportion of suprathreshold connections falling within the boundaries of each functional community defined in this parcellation were tracked along each gradient using line plots. Networks with proportions above chance level (1/7 networks = 0.14) were projected onto the cortical surface for each gradient anchor. (**Figure 3A**).

#### Functional organization along MPC gradients

Based on the thresholded and weighted functional connectivity matrix, we computed the participation coefficients (PC) of each vertex with respect to their functional network affiliation. This metric, as implemented in the brain connectivity toolbox (http://brain-connectivity-toolbox.net) (71), reflects the isotropy of connectivity of a given node to large-scale communities (72). Nodes with a more diverse pattern of connectivity to all modules of a network will tend to show higher participation coefficients (**Figure 3B**). We also applied a recently developed index of the average distance of functional connections (35, 73), also referred to as connectivity distance (CD). We used thresholded vertex-wise functional connectomes (keeping the top 10% – see *above*) and calculated the average geodesic distance to all other regions. This approach quantifies the average proximity of a given node’s strongest functional connections, and evaluates the balance of short- and long-range functional connections for cortical areas (**Figure 3C**). Spearman correlations assessed associations between these metrics (PC, CD) and each microstructural gradient, and statistical significance was determined via Moran spectral randomization (see *above*) (41). In addition, we computed four sets of standardized residuals from a simple linear model for PC and CD against each gradient independently. Spearman correlations were then computed between residual PC and CD and the microstructural gradient excluded from the linear model. This approach assessed the strength of the association between these metrics of functional organization and a specific axis of microstructural differentiation captured by each gradient while controlling for partial overlap between G1 and G2.

### Associations with fMRI meta-analytical activation maps

We examined which psychological processes were associated with changes resting-state functional connectivity profiles along both microstructural gradients, using meta-analytic term maps from the Neurosynth database (http://www.neurosynth.org) (44). This tool combines automated text mining and meta-analytical techniques to produce probabilistic mappings between cognitive terms and spatial brain response patterns. Z-statistic maps associated with 24 terms related to diverse domains (24, 34) were mapped to the cortical surface (FDR-corrected, *p* < 0.01). We retained only term maps encompassing the territory of the insula defined in the mask used in this study and mapping onto a minimum of 2% of vertices. G1 values were ranked, and retained terms were ordered according to the mean rank position along G1 (**Figure 4A**). To emphasize the unique ventral-dorsal transition captured by G2 and control for the partial overlap between both gradients, we repeated the same procedure on the rank differences between G2 and G1 (**Figure 4B**).

### Replication in independent sample

#### Sample characteristics

Our Replication sample (MICA-MTL cohort) consisted of 34 healthy adults (14 women; 29.38±6.04 years) recruited for high-resolution MRI that included quantitative T1 relaxometry as well as resting-state fMRI in our centre. Participants provided informed consent, and the study was approved by the research ethics board of the Montreal Neurological Institute and Hospital.

#### MRI acquisition

MRI data for the MICA-MTL cohort was acquired on a 3T Siemens Magnetom Prisma-Fit with a 64-channel head coil. Two T1w scans with identical parameters were acquired with a 3D-MPRAGE sequence (0.8mm isotropic voxels, matrix=320×320, 224 sagittal slices, TR=2300ms, TE=3.14ms, TI=900ms, flip angle=9°, iPAT=2). Quantitative T1 (qT1) relaxometry data was acquired using a 3D-MP2RAGE sequence (0.8mm isotropic voxels, 240 sagittal slices, TR=5000ms, TE=2.9ms, TI 1=940ms, T1 2=2830ms, flip angle 1=4°, flip angle 2=5°, iPAT=3, bandwidth=270 Hz/px, echo spacing=7.2ms, partial Fourier=6/8). We combined two inversion images for qT1 mapping in order to minimise sensitivity to B1 inhomogeneities and optimize intra- and intersubject reliability (74, 75).

#### Image preprocessing

Cortical surfaces were extracted from the T1w images using FreeSurfer 6.0 (68–70). Accuracy of surface extractions was visually inspected and corrected for any segmentation errors. 14 equivolumetric intracortical surfaces were then generated between the pial and white matter boundaries (32). qT1 volume was registered to Freesurfer native space using a boundary-based registration (59), and qT1 intensity values were sampled along the equivolumetric surfaces. We registered FreeSurfer native space to fsaverage5 space using Caret5 landmark-based registration (76) to resample the evaluated surfaces to a common space.

#### MPC gradient replication

We applied the same approach as in the HCP sample to construct microstructural gradients in the Replication cohort. Intracortical qT1 profiles of vertices in our previously defined insula mask were extracted prior to building MPC matrices per hemisphere. MPC matrices were computed from qT1 data, and subsequently converted to affinity matrices. These matrices were mapped into a lower dimensional manifold with diffusion map embedding applying identical parameters to those used in the Discovery cohort (33). Resulting gradients were aligned to those obtained with T1w/T2w data of the corresponding hemisphere using Procrustes analysis (63). Only the first two gradients were retained and correlated with microstructural gradients generated in the HCP sample using Spearman correlations.

### Code and data availability statement

Code used to generate the MPC and microstructural gradients, as well as Discovery and Replication sample microstructural gradient maps are available on https://github.com/MICA-MNI/micaopen.

## ACKNOWLEDGEMENTS

J.R. received support from the Canadian Open Neuroscience Platform (CONP). C.P. received support from the Transforming Autism Care Consortium (TACC) and Fonds de la Recherche du Québec – Santé (FRQ-S). S.L. acknowledges funding from FRQ-S and the Canadian Institutes of Health Research (CIHR). R.V.d.W. receives support from a Savoy Foundation studentship. B.F. receives salary support from FRQ-S (Chercheur-Boursier clinician Junior 2) and start-up funding from the Montreal Neurological Institute. B.C.B. acknowledges support from CIHR (FDN-154298), SickKids Foundation (NI17-039), Natural Sciences and Engineering Research Council (NSERC; Discovery-1304413), Azrieli Center for Autism Research of the Montreal Neurological Institute (ACAR), and salary support from FRQS (Chercheur Boursier Junior 1).

