## Supplementary Materials for "Myeloarchitecture gradients in the human insula serve as blueprints for its diverse connectivity and function"

**RUNNING TITLE**
Myeloarchitecture blueprints in the insula

**CORRESPONDING AUTHORS**

Boris C. Bernhardt, PhD
Multimodal Imaging and Connectome Analysis Lab
McConnell Brain Imaging Centre
Montreal Neurological Institute and Hospital
McGill University, Montreal, Quebec, H3A 2B4
Canada

Phone : (+1) 514 398 3044

Jessica Royer, PsyD
Multimodal Imaging and Connectome Analysis Lab
McConnell Brain Imaging Centre
Montreal Neurological Institute and Hospital
McGill University, Montreal, Quebec, H3A 2B4
Canada


**SUPPLEMENTARY FIGURE 1**

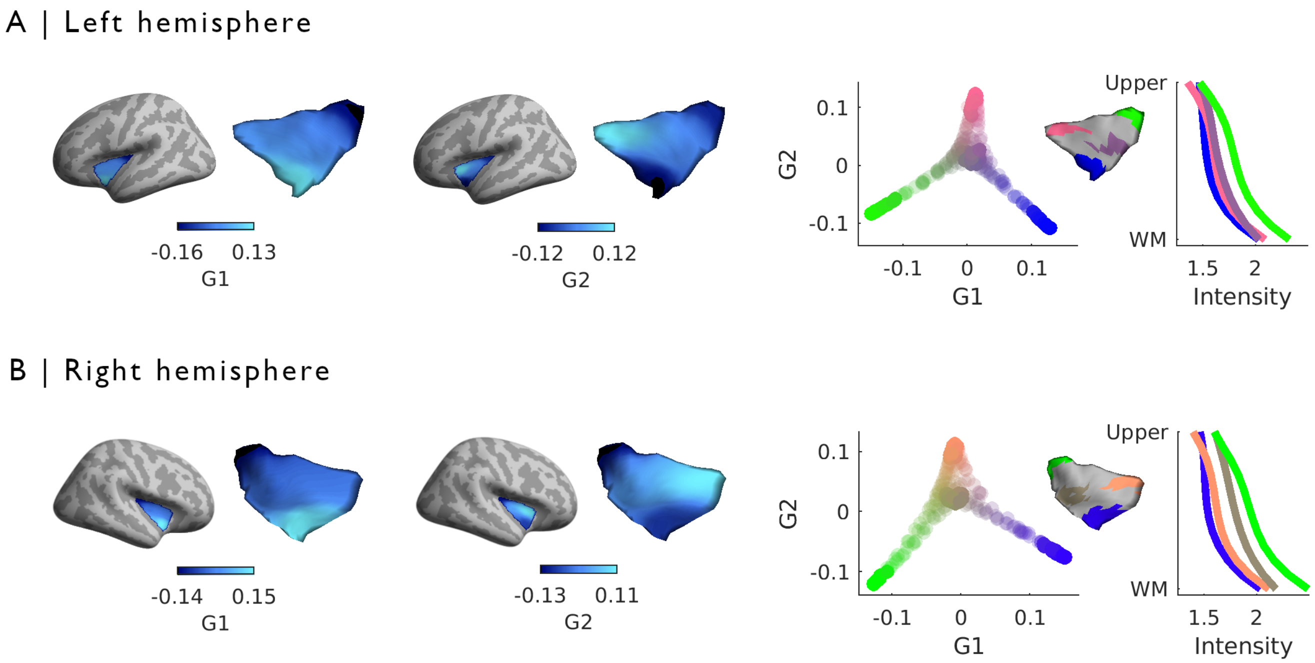


**Figure S1. In vivo microstructural gradients of the insular cortex are replicable across hemispheres**. Microstructural profile covariance (MPC) gradients in the left (A) and right (B; see also *Figure 1*) hemisphere were highly similar. The principal gradient G1 followed the anterior-posterior axis of the insula, differentiating posterior and ventral anterior regions. The second gradient G2 evolved along a dorsal-ventral pattern, and was anchored in dorsal anterior areas and radiated towards posterior and ventral anterior insular banks.

### SUPPLEMENTARY FIGURE 2

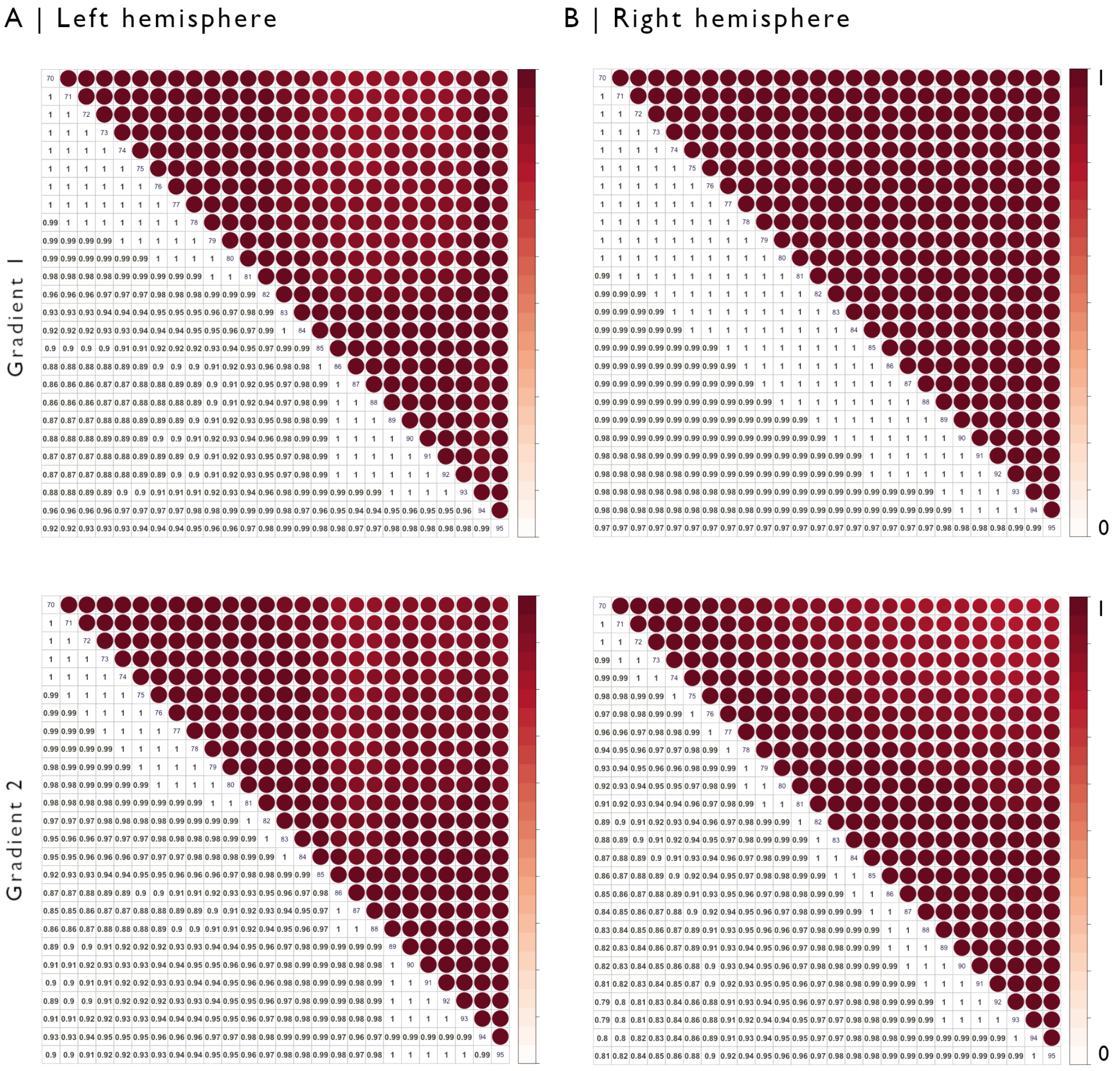


**Figure S2. Robustness to changes in MPC matrix sparsity.** Row-wise thresholding of the MPC matrix was systematically varied between 70 and 95% and effect on resulting gradients was assessed. Correlation matrices show Spearman correlation coefficients for all threshold combinations independently for each hemisphere (A, Left hemisphere ; B, Right hemisphere) and for the first two gradients (Top, G1 ; Bottom, G2).

**SUPPLEMENTARY FIGURE 3**

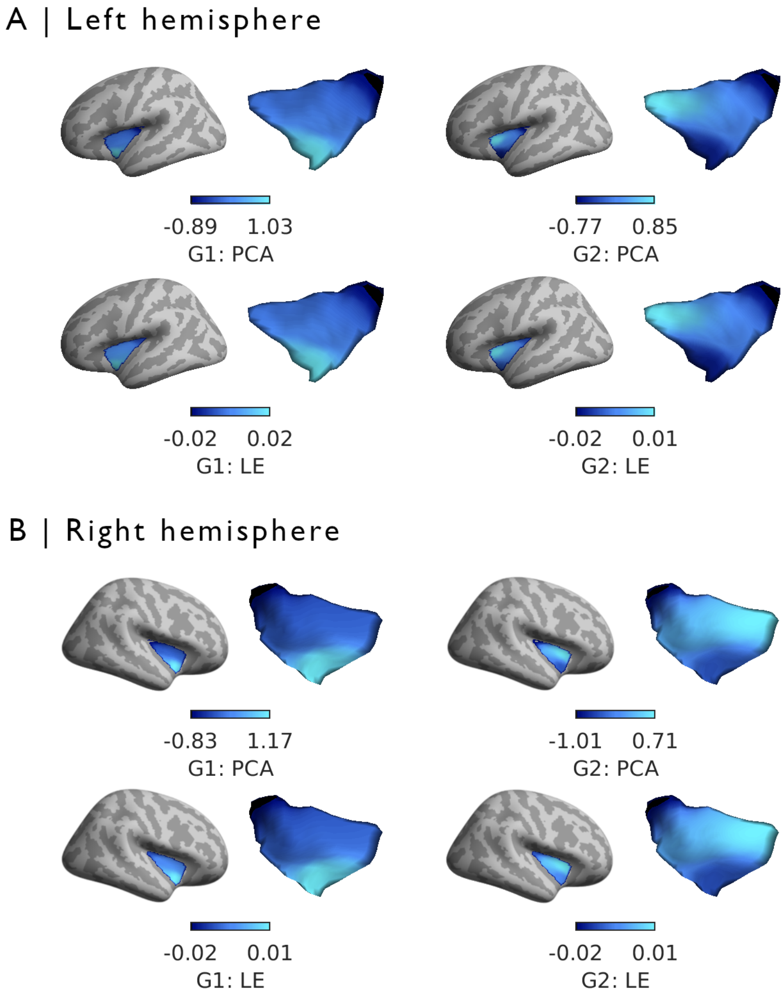


**Figure S3. Robustness to different dimensionality reduction techniques.** MPC gradients were generated from the thresholded MPC matrix (sparsity = 90) using different dimensionality reduction approaches. Results are presented for the left (A) and right (B) hemisphere independently using principal component analysis (PCA ; top), and laplacian eigenmaps (bottom ; LE). Correlations between presented gradient findings obtained with diffusion map embedding were strongly correlated with solutions yielded by PCA (Left : 0.98/0.97 G1/G2 ; Right : 0.99/0.99 G1/G2) and LE (Left : 0.99/0.98 G1/G2 ; Right : 0.99/0.99 G1/G2).

**SUPPLEMENTARY FIGURE 4**

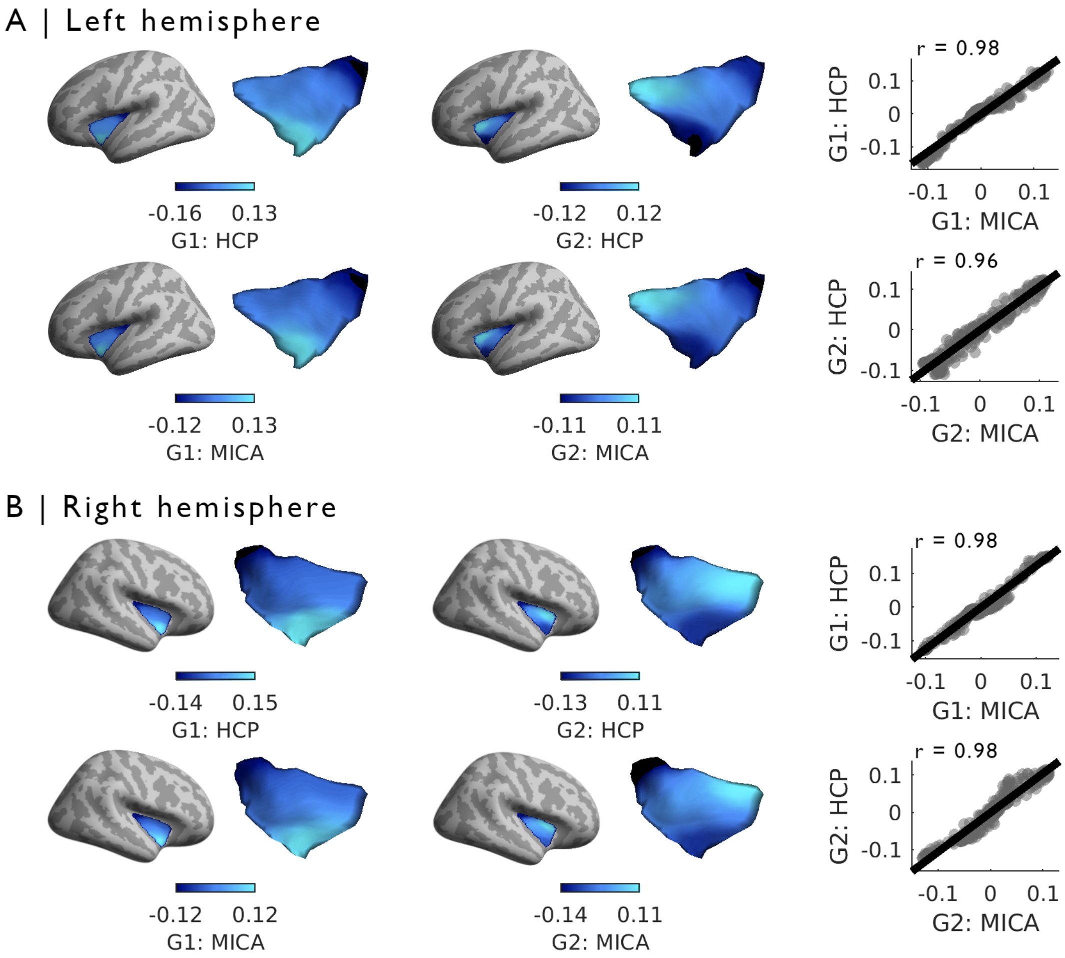


**Figure S4. Replication of microstructural gradients in an independent dataset. (A)** Left hemisphere microstructural gradients generated in the HCP sample with T1w/T2w image intensities were highly correlated with gradients obtained with qT1 imaging in the MICA-MTL sample after alignment using Procrustes analysis. **(B)** MPC gradient findings were also highly correlated across samples in the right hemisphere.
